# Genetic studies of accelerometer-based sleep measures in 85,670 individuals yield new insights into human sleep behaviour

**DOI:** 10.1101/303925

**Authors:** Samuel E. Jones, Vincent T. van Hees, Diego R. Mazzotti, Pedro Marques-Vidal, Séverine Sabia, Ashley van der Spek, Hassan S. Dashti, Jorgen Engmann, Desana Kocevska, Jessica Tyrrell, Robin N. Beaumont, Melvyn Hillsdon, Katherine S. Ruth, Marcus A. Tuke, Hanieh Yaghootkar, Seth Sharp, Yingjie Jie, Jamie W. Harrison, Rachel M. Freathy, Anna Murray, Annemarie I. Luik, Najaf Amin, Jacqueline M. Lane, Richa Saxena, Martin K. Rutter, Henning Tiemeier, Zoltan Kutalik, Meena Kumari, Timothy M. Frayling, Michael N. Weedon, Philip Gehrman, Andrew R. Wood

## Abstract

Sleep is an essential human function but its regulation is poorly understood. Identifying genetic variants associated with quality, quantity and timing of sleep will provide biological insights into the regulation of sleep and potential links with disease. Using accelerometer data from 85,670 individuals in the UK Biobank, we performed a genome-wide association study of 8 accelerometer-derived sleep traits, 5 of which are not accessible through self-report alone. We identified 47 genetic associations across the sleep traits (*P*<5×10^-8^) and replicated our findings in 5,819 individuals from 3 independent studies. These included 26 novel associations for sleep quality and 10 for nocturnal sleep duration. The majority of newly identified variants were associated with a single sleep trait, except for variants previously associated with restless legs syndrome that were associated with multiple sleep traits. Of the new associated and replicated sleep duration loci, we were able to fine-map a missense variant (p.Tyr727Cys) in *PDE11A*, a dual-specificity 3’,5’-cyclic nucleotide phosphodiesterase expressed in the hippocampus, as the likely causal variant. As a group, sleep quality loci were enriched for serotonin processing genes and all sleep traits were enriched for cerebellar-expressed genes. These findings provide new biological insights into sleep characteristics.

## INTRODUCTION

Sleep is an essential human function, but many aspects of its regulation remain poorly understood. Adequate sleep is important for health and wellbeing, and changes in sleep quality, quantity and timing are strongly associated with several human diseases and psychiatric disorders^1-5^. Identifying genetic variants influencing sleep traits will provide new insights into the molecular regulation of sleep in humans and help to establish the genetic contribution to causal links between sleep and associated chronic diseases, such as diabetes and obesity^6-10^.

Genome-wide association studies (GWAS) are an important first step towards the discovery of new biological mechanisms of complex traits. Previous large-scale genetic studies of sleep traits have relied on self-reported measures. For example, using questionnaire data from 47,180 individuals, the CHARGE consortium identified the first common genetic variant, near *PAX8*, robustly associated with sleep duration^11^. Subsequent studies in up to 128,286 individuals using the interim data release of the UK Biobank identified two additional sleep duration loci^12^,^13^ and a parallel analysis of the full UK Biobank release of 446,118 individuals identified a total of 78 associated loci (Dashti *et al*., BioRxiv 2018, https://doi.org/10.1101/274977). Genetic associations have also been identified for other self-reported sleep traits including chronotype^12^,^14^,^15^, insomnia, and daytime sleepiness^13^,^16-18^ (Jansen *et al.* BioRxiv 2018, https://doi.org/10.1101/214973 and Lane *et al.* BioRxiv 2018: https://doi.org/10.1101/257956).

Although the reported associations revealed relevant pathways related to mechanisms underlying sleep regulation, in large scale studies self-report measures are typically based on a limited number of questions that only approximate a limited number of sleep traits and may be subject to bias related to an individual’s perception and recall of sleeping patterns^19-23^. Polysomnography (PSG) is regarded as the “gold standard” method of quantifying nocturnal sleep traits, but it is impractical to perform in large cohorts. Additionally, PSG is relatively burdensome for the participant making it less suitable for measuring sleep over multiple nights and capturing inter-daily variability. Research-grade activity monitors (accelerometers), also known as actigraphy devices, provide cost-effective estimates of sleep using validated algorithms^24^,^25^. However, accelerometer-based studies have often involved much smaller sample sizes than those required for GWAS and have generally focussed on day-time activity^26^,^27^. The UK Biobank study is a unique resource collecting vast amounts of clinical, biomarker, and questionnaire data on ∼500,000 UK residents. Of these, 103,000 participants wore activity monitors continuously for up to 7 days. This provides an unprecedented opportunity to derive accelerometer-based estimates of sleep quality, quantity and timing and to assess the genetics of sleep traits.

In this study we identify genetic variants associated with objective measures of sleep and rest-activity patterns and use them to further understand the biology of sleep. We used accelerometer data from the UK Biobank to extract estimates of sleep characteristics using a heuristic method previously validated using independent PSG datasets^28^,^29^. We analysed a total of 8 accelerometer-based measures of sleep quality (sleep efficiency and the number of nocturnal sleep episodes), timing (sleep-midpoint, timing of the least active 5 hours (L5), and timing of the most active 10 hours (M10)), and duration (diurnal inactivity and nocturnal sleep duration and variability) by performing a GWAS in 85,670 UK Biobank participants and assess replication of the findings in 3 independent studies. Our analysis primarily focuses on traits that cannot be captured, or are unavailable, from self-report sleep measures, and are likely to be underpowered for GWAS in studies with PSG data due to limited sample sizes.

## RESULTS

### Measures of sleep quality and quantity are not correlated with sleep timing

Descriptive statistics and correlations between the eight accelerometer-derived phenotypes are shown in **Supplementary Tables 1** and **2**. We observed little observational correlation (*R*) between measures of sleep timing and measures of nocturnal sleep duration and quality (−0.10 ≤ *R* ≤ 0.12). These negligible or limited correlations between timing and duration are consistent with data from chronotype and self-report sleep duration measures (*R* = −0.01). We also observed limited correlation between sleep duration and sleep quality as represented by the number of nocturnal sleep episodes (*R* = 0.14) but observed a stronger correlation between sleep duration and sleep efficiency (*R*= 0.57). The correlations between self-reported sleep duration and accelerometer-derived sleep duration was 0.19 and between self-reported chronotype (“morningness”) and L5 timing was −0.29.

### Accelerometer-derived estimates of sleep patterns are heritable

To estimate the proportion of variance attributable to genetic factors for a given trait, we used BOLT-REML to estimate SNP-based heritability (*h*^*2*^_*SNP*_) (**Table 1**). *h*^*2*^_*SNP*_ estimates ranged from 2.8% (95% CI 2.0%, 3.6%) for variation in sleep duration (defined as the standard deviation of accelerometer-derived sleep duration across all nights), to 22.3% (95% CI 21.5%, 23.1%) for number of nocturnal sleep episodes. For sleep duration, we observed higher heritability using the accelerometer-derived measure (*h*^*2*^_*SNP*_= 19.0%, 95% CI 18.2%, 19.8%) in comparison to self-report sleep duration (*h* ^*2*^ *SNP* = 8.8%, 95% CI 8.6%, 9.0%). The heritability estimates for sleep and activity timings (maximum *h*^*2*^ *SNP*= 11.7%, 95% CI 10.9%, 12.5%) were lower than for self-report chronotype (*h*^*2*^_*SNP*_ = 13.7%, 95% CI 13.3%, 14.0%) (Jones *et al*. BioRxiv 2018, https://doi.org/10.1101/303941).

**Table 1.** Pseudo-heritability estimates of derived sleep variables from BOLT-REML

| **Sleep variable** | **$h_2$** | **95% CI** |
| --- | --- | --- |
| Sleep duration | 0.190 | 0.182 – 0.198 |
| Sleep duration variability (SD) | 0.028 | 0.020 – 0.036 |
| Number of nocturnal sleep episodes | 0.223 | 0.215 – 0.231 |
| Sleep efficiency | 0.130 | 0.122 – 0.138 |
| L5 timing | 0.117 | 0.109 – 0.125 |
| M10 timing | 0.087 | 0.079 – 0.095 |
| Sleep midpoint timing | 0.101 | 0.093 – 0.109 |
| Diurnal Inactivity | 0.148 | 0.134 – 0.161 |

### Low genetic correlation between self-reported and accelerometer-derived sleep duration

To quantify the genetic contribution, overlap between accelerometer-derived and self-reported sleep traits, we performed genetic correlation analyses using LD-score regression as implemented in LD-Hub^30^. We observed strong genetic correlations between L5, M10 and sleep midpoint timing and chronotype (r_G_>0.79) and weaker genetic correlation between objective versus self-reported sleep duration (r_G_=0.43). This observation suggests differences in the genetic contribution to variation in self-reported versus objective sleep duration.

### Forty-seven genetic associations identified across the accelerometer-derived sleep traits

To identify genetic loci associated with accelerometer-derived sleep traits, we performed a genome-wide association analysis of 11,977,111 variants in up to 85,670 individuals for the 8 accelerometer-derived sleep traits. We identified 47 genetic associations across 7 of the phenotypes at the standard GWAS threshold (*P*<5×10^-8^). Among these associations, 20 reached a more stringent threshold of *P*<8×10^-10^. We estimate that this threshold reflects a better type 1 error rate to account for the approximate number of independent genetic variants analysed (Jones *et al*. BioRxiv 2018, https://doi.org/10.1101/303941) against all 8 accelerometer-based traits (**Table 2** and **Supplementary Figs 1-2**). Twenty-six associations were observed for sleep quality measures, including 21 variants associated with number of nocturnal sleep episodes and 5 associated with sleep efficiency (8 and 2 at *P*<8×10^-10^, respectively). An additional 8 genetic associations were identified for sleep and activity timing. These included 6 associated with L5 timing, 1 associated with M10 timing, and 1 associated with mid-point sleep. Only 3 associations with L5 timing were detected at *P*<8×10^-10^. Finally, for sleep duration we observed 13 associations – 11 for sleep duration and 2 associated with diurnal inactivity (6 and 1 at *P*<8×10^-10^, respectively). Of these 47 associations reaching *P*<5×10^-8^ and the 20 associations reaching *P*<8×10^-10^, 31 and 9 were not previously reported in studies based on self-report measures, respectively (**Table 2**). The variance explained by all the discovered loci ranged from 0.04% for sleep midpoint timing to 0.8% for number of nocturnal sleep episodes. The lambda GC observed across these analyses ranged from 1.03 (sleep duration variability) to 1.14 (number of nocturnal sleep episodes), while LD-Score intercepts ranged from 1.03 (diurnal inactivity) to 1.07 (sleep midpoint timing). These results suggest that any inflation of test statistics observed is more likely to due to the polygenicity of the phenotype tested over and above population stratification.

**Table 2.** Summary statistics for 47 genetic associations identified in the UK Biobank reaching P<5×10^-8^

| TRAIT | SNP | Chr | BP (hg19) | E/O<br>Allele | EA<br>Freq | BETA | SE | P | Gene Region |
| --- | --- | --- | --- | --- | --- | --- | --- | --- | --- |
| L5 timing | rs1144566 | 1 | 182,569,626 | C/T | 0.970 | 0.096 | 0.014 | 8E-12 | *RGS16/RNASEL* |
| L5 timing | rs113851554 | 2 | 66,750,564 | T/G | 0.057 | 0.133 | 0.011 | 2E-35 | *MEIS1\** |
| L5 timing | rs12991815 | 2 | 68,071,990 | C/G | 0.424 | 0.029 | 0.005 | 2E-09 | *C1D\** |
| L5 timing | rs9369062 | 6 | 38,437,303 | A/C | 0.708 | 0.039 | 0.005 | 9E-14 | *BTBD9\** |
| L5 timing | rs4882315 | 12 | 38,458,906 | T/C | 0.507 | 0.027 | 0.005 | 2E-08 | *CPNE8/ALG10B* |
| L5 timing | rs12927162 | 16 | 52,684,916 | G/A | 0.277 | 0.029 | 0.005 | 3E-08 | *TOX3\** |
| M10 timing | rs1973293 | 12 | 38,679,575 | C/T | 0.481 | 0.029 | 0.005 | 1E-09 | *CPNE8/ALG10B* |
| Sleep duration | rs2660302 | 1 | 98,520,219 | A/T | 0.811 | 0.041 | 0.006 | 9E-12 | *DPYD* |
| Sleep duration | rs113851554 | 2 | 66,750,564 | G/T | 0.943 | 0.110 | 0.011 | 2E-25 | *MEIS1\** |
| Sleep duration | rs62158170 | 2 | 114,082,175 | G/A | 0.217 | 0.054 | 0.006 | 3E-21 | *PAX8* |
| Sleep duration | rs17400325 | 2 | 178,565,913 | T/C | 0.958 | 0.066 | 0.012 | 2E-08 | *PDE11A* |
| Sleep duration | rs72828540 | 6 | 19,102,286 | T/C | 0.752 | 0.041 | 0.005 | 1E-13 | *LOC101928519* |
| Sleep duration | rs9369062 | 6 | 38,437,303 | C/A | 0.292 | 0.033 | 0.005 | 2E-10 | *BTBD9\** |
| Sleep duration | rs2975734 | 8 | 10,090,097 | C/G | 0.561 | 0.027 | 0.005 | 1E-08 | *MSRA* |
| Sleep duration | rs13282541 | 8 | 41,723,550 | C/T | 0.739 | 0.032 | 0.005 | 4E-09 | *ANK1* |
| Sleep duration | rs2880370 | 8 | 105,987,057 | A/T | 0.670 | 0.028 | 0.005 | 2E-08 | *LRP12/ZFPM2* |
| Sleep duration | rs800165 | 12 | 67,645,219 | C/T | 0.343 | 0.028 | 0.005 | 3E-08 | *CAND1* |
| Sleep duration | rs10138240 | 14 | 63,353,479 | G/C | 0.514 | 0.029 | 0.005 | 7E-10 | *KCNH5* |
| Sleep midpoint | rs11892220 | 2 | 231,691,067 | T/A | 0.339 | 0.029 | 0.005 | 3E-08 | *CAB39* |
| Sleep efficiency | rs113851554 | 2 | 66,750,564 | G/T | 0.943 | 0.101 | 0.011 | 5E-22 | *MEIS1\** |
| Sleep efficiency | rs62158169 | 2 | 114,081,827 | T/C | 0.216 | 0.032 | 0.006 | 2E-08 | *PAX8* |
| Sleep efficiency | rs17400325 | 2 | 178,565,913 | T/C | 0.958 | 0.074 | 0.012 | 2E-10 | *PDE11A* |
| Sleep efficiency | rs13094687 | 3 | 52,450,043 | G/A | 0.315 | 0.029 | 0.005 | 1E-08 | *PHF7* |
| Sleep efficiency | rs13080973 | 3 | 138,596,050 | G/A | 0.202 | 0.032 | 0.006 | 3E-08 | *FOXL2* |
| No. sleep episodes | rs12714404 | 2 | 282,462 | T/G | 0.283 | 0.037 | 0.005 | 1E-12 | *ACP1/SH3YL1* |
| No. sleep episodes | rs310727 | 3 | 4,336,589 | T/C | 0.475 | 0.026 | 0.005 | 3E-08 | *SUMF1/SETMAR* |
| No. sleep episodes | rs55754932 | 3 | 87,847,754 | C/A | 0.284 | 0.037 | 0.005 | 2E-12 | *HTR1F* |
| No. sleep episodes | rs9864672 | 3 | 137,076,353 | T/C | 0.522 | 0.029 | 0.005 | 2E-10 | *IL20RB/SOX14* |
| No. sleep episodes | rs4974697 | 4 | 2,473,092 | T/A | 0.390 | 0.026 | 0.005 | 5E-08 | *RNF4* |
| No. sleep episodes | rs7377083 | 4 | 102,708,997 | A/C | 0.430 | 0.029 | 0.005 | 2E-09 | *BANK1* |
| No. sleep episodes | rs749100 | 5 | 63,307,862 | A/G | 0.582 | 0.033 | 0.005 | 9E-12 | *HTR1A/RNF180* |
| No. sleep episodes | rs9341399 | 6 | 73,773,644 | C/T | 0.936 | 0.066 | 0.010 | 6E-12 | *KCNQ5* |
| No. sleep episodes | rs1889978 | 6 | 124,771,233 | C/T | 0.485 | 0.027 | 0.005 | 5E-09 | *NKAIN2* |
| No. sleep episodes | rs2141277 | 7 | 39,099,178 | A/G | 0.478 | 0.026 | 0.005 | 1E-08 | *POU6F2* |
| No. sleep episodes | rs10233848 | 7 | 103,122,645 | G/A | 0.293 | 0.035 | 0.005 | 2E-11 | *RELN* |
| No. sleep episodes | rs1124116 | 10 | 99,371,147 | A/G | 0.730 | 0.031 | 0.005 | 2E-09 | *HOGA1/MORN4* |
| No. sleep episodes | rs4755731 | 11 | 43,685,168 | G/A | 0.431 | 0.028 | 0.005 | 3E-09 | *HSD17B12* |
| No. sleep episodes | rs3751837 | 16 | 3,583,173 | C/T | 0.781 | 0.033 | 0.006 | 4E-09 | *CLUAP1* |
| No. sleep episodes | rs8045740 | 16 | 20,262,776 | G/T | 0.868 | 0.052 | 0.007 | 6E-14 | *GPR139* |
| No. sleep episodes | rs11078917 | 17 | 37,746,359 | A/C | 0.279 | 0.029 | 0.005 | 3E-08 | *NEUROD2* |
| No. sleep episodes | rs11082030 | 18 | 35,501,739 | T/C | 0.725 | 0.030 | 0.005 | 8E-09 | *CELF4* |
| No. sleep episodes | rs8098424 | 18 | 52,458,218 | G/A | 0.619 | 0.027 | 0.005 | 1E-08 | *RAB27B* |
| No. sleep episodes | rs76753486 | 19 | 42,684,264 | T/C | 0.084 | 0.047 | 0.008 | 2E-08 | *DEDD2/ZNF526* |
| No. sleep episodes | rs429358 | 19 | 45,411,941 | T/C | 0.848 | 0.036 | 0.007 | 4E-08 | *APOE* |
| No. sleep episodes | rs12479469 | 20 | 61,145,196 | A/G | 0.342 | 0.031 | 0.005 | 4E-10 | *MIR133A2* |
| Diurnal inactivity | rs17805200 | 9 | 13,764,434 | C/T | 0.272 | 0.031 | 0.005 | 5E-09 | *MPDZ/NFIB* |
| Diurnal inactivity | rs7155227 | 14 | 63,365,094 | T/G | 0.523 | 0.033 | 0.005 | 2E-12 | *KCNH5* |

CHR=chromosome; BP=base-pair position (GRCh37/hg19); EA/OA=effect allele/other allele, EA Freq=effect allele frequency; SE=standard error; L5 timing=midpoint of least active 5 hours; M10 timing=midpoint of most active 10 hours; No. sleep episodes=number of nocturnal sleep episodes; \* locus previously reported for Restless Legs Syndrome<sup>31</sup>.

**Figure 1.**
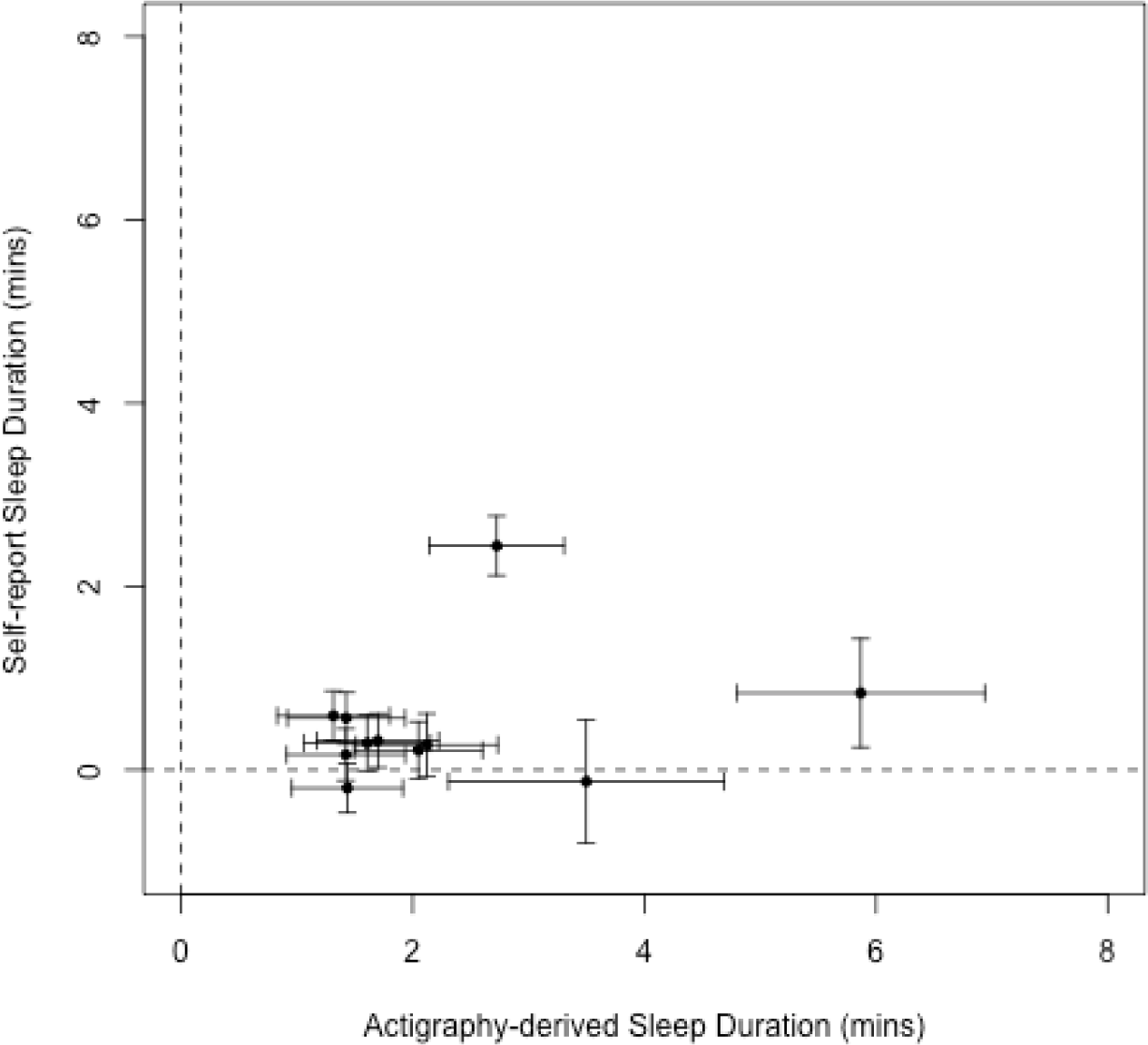
Comparisons of betas for 11 genetic variants associated with accelerometer-derived sleep duration against betas from a parallel GWAS of self-report sleep duration.

**Figure 2.**
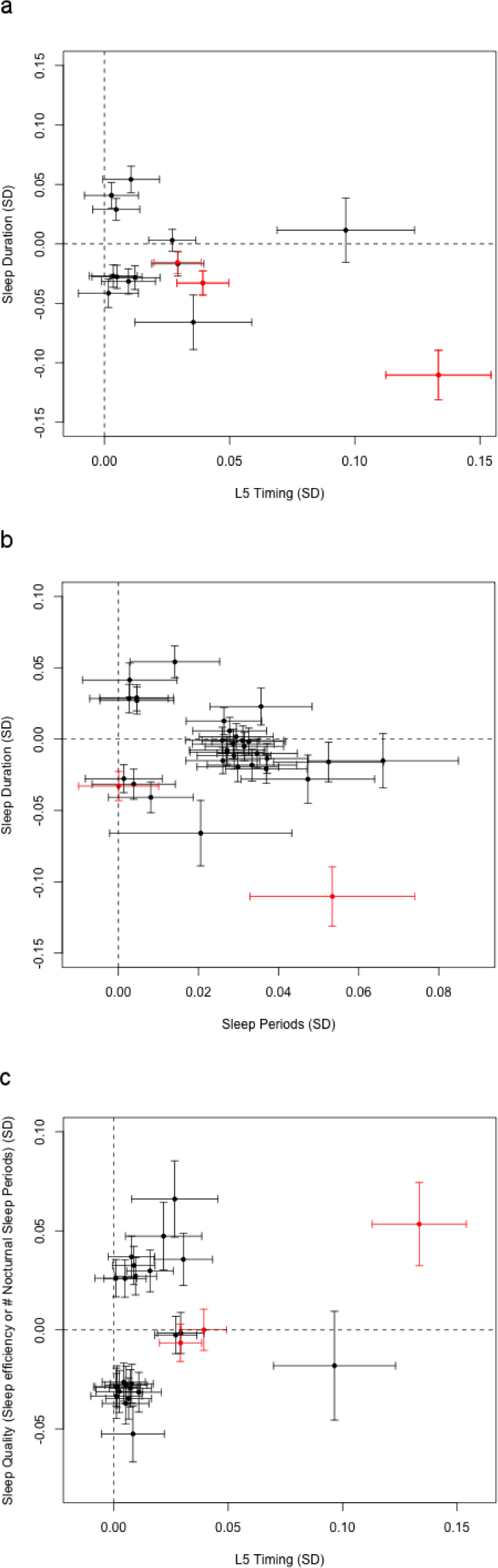
Comparison of betas for genetic variants associated with **a)** either L5 timing or sleep duration, **b)** either sleep duration or the number of nocturnal sleep episodes, and **c)** either L5 timing or sleep quality (number of nocturnal sleep episodes or sleep efficiency). Variants previously associated with restless legs syndrome are highlighted in red. Betas represent standard deviations of the inverse-normal distribution of each trait.

### Replication of 47 genetic associations in 5,819 individuals

We attempted to replicate our findings in up to 5,819 adults from the Whitehall (N=2,144), CoLaus (N=2,257), and Rotterdam Study (subsample from RS-I, RS-II and RS-III, N=1,418) who had worn similar wrist-worn tri-axial accelerometer devices for a comparable duration as the UK Biobank participants. Individual study and meta-analysis results for the three replication studies are presented in **Supplementary Table 3**. Of the 20 associations reaching *P*<8×10^-10^, 18 were directionally consistent in the replication cohort meta-analyses (*P*_*binomial*_ = 3×10^-4^). Of the additional 27 signals, 18 were directionally consistent in the replication meta-analysis (*P*_*binomial*_ = 0.03). For traits with more than one SNP associated at *P*<5×10^-8^ in the UK Biobank, we combined the effects of each SNP (aligned to the trait increasing allele) and tested them in the replication data. In the combined effects analysis, we observed overall associations with sleep duration (*P*=0.008), sleep efficiency (*P*=3×10^-4^), number of nocturnal sleep episodes (*P*=2×10^-6^), and sleep timing (*P*=0.034) (**Supplementary Tables 3** and **4**).

### Variants associated with sleep quality include known restless legs syndrome, sleep duration, and cognitive decline associated variants

Of the 5 variants associated with sleep efficiency, one was the strongly associated *PAX8* sleep duration signal^11^ and one was a restless legs syndrome/insomnia associated signal (*MEIS1*)^17^,^31^. Of the 20 loci associated with number of nocturnal sleep episodes, one is represented by the *APOE* variant (rs429358). This variant is a proxy for the *APOE* ε*4* risk allele that is strongly associated with late-onset Alzheimer’s disease and cognitive decline^32^. The ε*4* allele is associated with a reduced number of nocturnal sleep episodes (−0.13 sleep episodes; 95% CI: −0.16, - 0.11; *P*=4×10^-8^). This finding is strengthened by additional analyses of the ε*2,* ε*3* and ε*4 APOE* Alzheimer’s disease risk alleles, with an overall reduction in the number of nocturnal sleep episodes observed with higher risk haplotypes (F(5, 72578)=5.36, *P*=0.001) (**Supplementary Table 5**). This finding is inconsistent with the observational association between cognitive decline in older age and poorer sleep quality^33-36^. We also noted that the APOE ε*4* risk allele was nominally associated (*P*<0.05) with sleep timing (L5, −1.8 minutes per allele, *P*=4×10^-6^; sleep-midpoint (−0.6 minutes per allele; *P*=0.002), sleep duration (−1.1 minutes per allele, *P*=7×10^-4^), and diurnal inactivity (−1.0 minutes per allele, *P*=2×10^-5^). Apart from the *APOE* variant (rs429358), which had double the effect size in the older half of the cohort (**Supplementary Table 5**), there were minimal differences in effect sizes in a range of sensitivity analyses, including removing individuals on sleep or depression medication, adjustments for BMI and lifestyle factors, and splitting the cohort by median age (**Supplementary Table 6** and **Supplementary Methods**).

### Six association signals identified for accelerometer-derived measures of sleep timing

We identified 6 loci associated with L5 timing, of which 3 have not previously been associated with self-report chronotype but have been associated with restless legs syndrome^31^. The index variants at these 3 loci are in strong to modest LD with the previously reported variants associated with restless legs syndrome (rs113851554, *MEIS1*, LD *r*^2^ = 1.00; rs12991815, *C1D*, LD *r*^2^ = 0.96; rs9369062, *BTBD9,* LD *r*^2^ = 0.49). The three variants that reside in loci previously associated with self-report chronotype are in strong to modest linkage disequilibrium with those previously reported^12^,^14^,^15^ (rs1144566, *RSG16*, LD *r*^2^ > 0.91; rs12927162 *TOX3,* LD *r*^2^ = 1.00; rs4882315, *ALG10B*, LD *r*^2^ = 0.58). The variant rs1144566 is a missense coding change (p.His137Arg) in exon 5 of *RSG16,* a known circadian rhythm gene which contains the variants strongly associated with self-report chronotype^12^. In a parallel self-report chronotype study in the UK Biobank, rs1144566 represented the strongest association, with the T allele having a morningness odds ratio of 1.26 (*P*=2×10^-95^) (Jones *et al.* BioRxiv 2018, https://doi.org/10.1101/303925). In addition, variants in the region of *TOX3* have previously been associated with restless legs syndrome^31^. However, our lead SNP (rs12927162) was not in LD with the previously reported index variant at this locus (rs45544231, LD *r*^*2*^ = 0.004). There were minimal differences in effect sizes when we performed a range of sensitivity analyses, including removing individuals on depression medication, adjustments for BMI and lifestyle factors and splitting the cohort by median age (**Supplementary Table 6** and **Supplementary Methods**).

### Ten novel sleep duration loci identified from accelerometer-derived sleep duration GWAS

We identified 11 loci associated with accelerometer-derived sleep duration, including ten not previously reported to be associated with self-report sleep duration, despite the 5-fold increase in sample size available for a parallel self-report sleep duration GWAS study (Dashti *et al.* BioRxiv 2018, https://doi.org/10.1101/274977; **Figure 1** and **Supplementary Table 7**). This lower overlap in signals is consistent with the lower genetic correlation between self-reported and objective sleep duration than between chronotype and objective measures of sleep and activity timing. The lead variants representing the ten new sleep duration loci all had the same direction and larger effects in the accelerometer data compared to self-report data, with effect sizes ranging from 1.3 to 5.9 minutes compared to 0.1 to 0.8 minutes (self-report *P*<0.05), with the *MEIS1* locus having the strongest effect. Two of the ten new sleep duration signals (rs113851554 in *MEIS1* and rs9369062 in *BTBD9*) have previously been associated with restless legs syndrome. The one variant previously detected based on self-report sleep duration, near *PAX8,* was the first variant to be associated with sleep duration through GWAS^11^. The minor *PAX8* allele effect size was consistent across accelerometer-derived measures of sleep duration (2.7 minutes per allele, 95% CI: 2.1 to 3.3, *P*=3×10^-21^) and self-report sleep duration (2.4 minutes per allele, 95% CI: 2.1 to 2.8, *P*=7×10^-49^). We observed similar effect sizes in a subset of 72,510 unrelated Europeans from the UK Biobank, when removing individuals on depression medication and after adjusting for BMI and lifestyle factors. To confirm that associations were not influenced by age-related differences in sleep, we confirmed that there was also no difference in effect sizes between younger and older individuals (above and below the median age of 63.7 years) (**Supplementary Table 6**).

### Fine-mapping analysis identifies multiple likely causal variants

To identify credible SNP sets likely to contain causal variants within 500Kb of lead SNPs (log_10_ Bayes Factor > 2) for each trait with a genetic association (*P*<5×10^-8^) we used FINEMAP^37^ (**Supplementary Table 8)**. Two loci contained a coding variant with a probability greater than 80% for being the causal variant. The first variant (rs17400325, MAF = 4.2%) was a missense variant (p.Tyr727Cys) in *PDE11A*, a phosphodiesterase highly expressed in the hippocampus that was associated with sleep duration and sleep efficiency. The other was the missense *APOE* variant representing the e4 allele, known to predispose to Alzheimer’s disease, which was associated with the number of nocturnal sleep episodes. Of the remaining loci, 5 fine-mapped variants are eQTLs in the Genotype-Tissue Expression (GTEx) project^38^. Of these only the fine-mapped variant at the *CLUAP1* locus was the lead variant for the corresponding eQTL (**Supplementary Table 8**). *CLUAP1* is a gene previous associated with photoreceptor maintenance that is associated with number of nocturnal sleep episodes^39^.

### Associated loci are enriched for genes expressed in the cerebellum and serotonin pathway-related genes

We used MAGMA^40^ to assess tissue enrichment of genes at associated loci across the sleep traits. All traits showed an enrichment of genes in the cerebellum (**Supplementary Figures 3 and 4**). Loci associated with number of nocturnal sleep episodes were enriched for genes involved in serotonin pathways (*P*_*Bonferroni*_=0.0003) (**Supplementary Table 9**).

**Figure 3.**
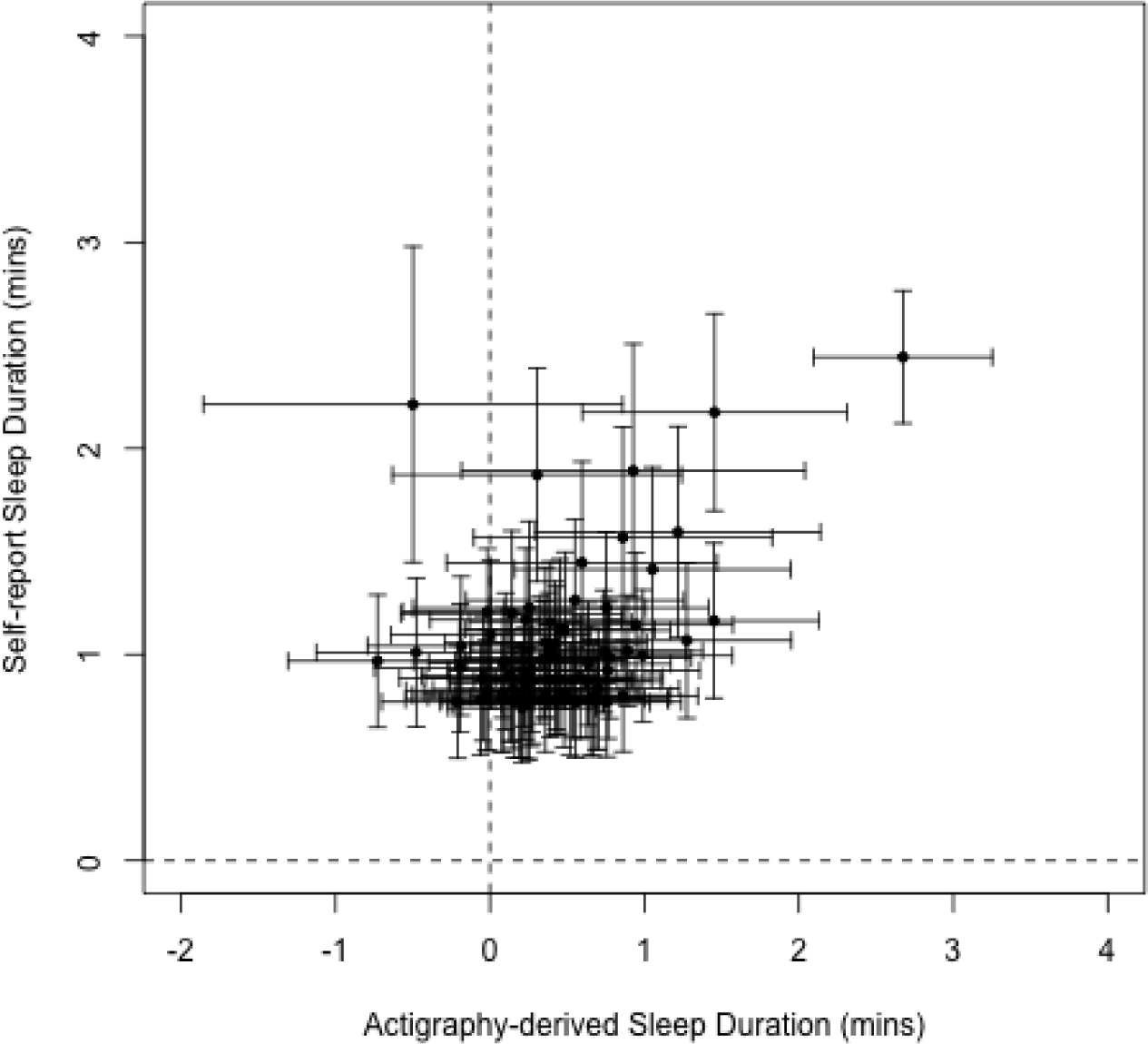
Comparisons of betas for 78 genetic variants associated with self-report sleep duration in a parallel GWAS effort (Dashti *et al,* BioRvix, 2018, https://doi.org/10.1101/274977).

**Figure 4.**
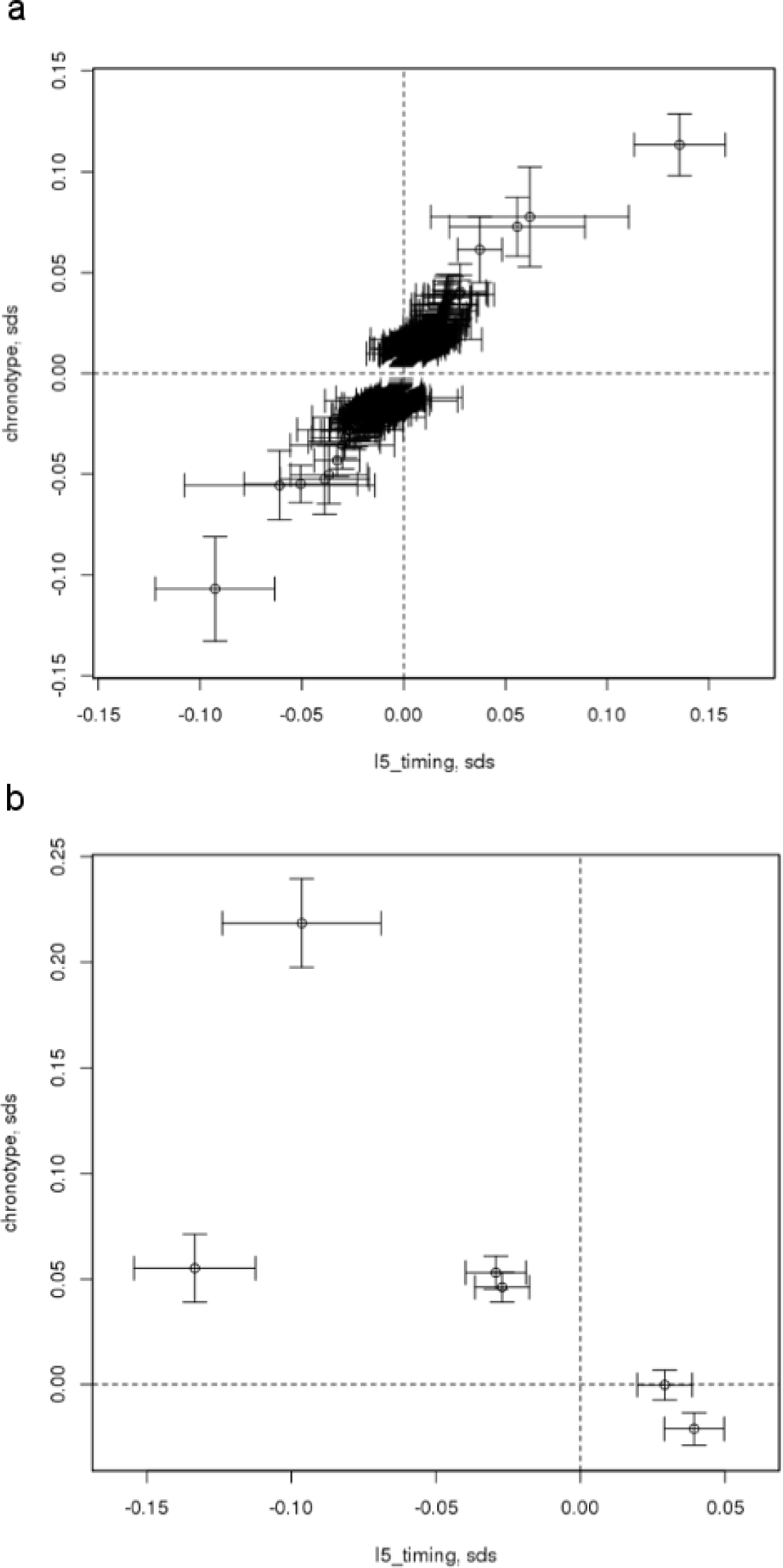
Correlations of genetic effect estimates based on self-report chronotype meta-analysis versus activity monitor midpoint sleep estimated from L5 timing for **a**) 351 variants identified from self-report chronotype GWAS and **b**) 6 variants identified for L5 timing from accelerometer derived estimates

### Multiple sleep traits have genetic variants previously associated with restless legs syndrome

We observed most variants to be associated with either sleep quality, duration, or timing, but not combinations of these sleep characteristics. However, the variant rs113851554 at the *MEIS1* locus was associated with sleep quality (sleep efficiency), duration, and timing (L5). In addition, the variant rs9369062 at the *BTBD9* locus was associated with both sleep duration and L5 timing. Both variants have previously been reported as associated with restless legs syndrome (**Figure 2**). To follow up this observation, we performed Mendelian Randomisation using 20 variants associated with restless legs syndrome in the discovery stage of the most recent and largest genome-wide association study^31^. We tested these 20 variants against all 8 activity-monitor derived sleep traits and showed a clear causative association of restless legs syndrome with all sleep traits. We also observed a causative association of restless legs syndrome with self-report sleep duration and chronotype, suggesting that variants associated with restless legs syndrome were not artefacts of the accelerometer-derived measures of sleep (**Supplementary Table 10**).

### Waist-hip-ratio (adjusted for BMI) and educational attainment causally influence sleep outcomes

Given genetic correlations are generally similar to observational correlations^41^, we used genetic correlations to prioritise traits for subsequent Mendelian Randomisation analyses. Using LD-Hub^30^ we tested for genetic correlation between the 8 activity monitor derived measures and 234 published GWAS studies across a range of diseases and traits. After adjustment for the number of genetic correlations tested (8 x 234), we observed genetic correlations between sleep traits and obesity and educational attainment related traits (**Supplementary Table 11**). After adjusting for the number of tests in the bi-directional MR analysis (99), we observed evidence that higher waist-hip-ratio (adjusted for BMI) is causally associated with lower sleep duration (*P*_*IVW*_ = 5×10^-6^) and lower sleep efficiency (*P*_*IVW*_ = 3×10^-4^). In addition, we observed higher educational attainment to be causally associated with lower sleep duration (*P*_*IVW*_ = 5×10^-5^) (**Supplementary Table 12**). We observed no evidence of causal effects of accelerometer-based sleep traits on outcomes tested (**Supplementary Table 13**).

### Estimates of the effects on accelerometer-derived and self-report-derived sleep traits are correlated

We compared effects of variants associated with self-reported sleep duration and chronotype identified in parallel GWAS analyses. Overall, we observed directional consistency with the accelerometer-derived measures. In a parallel GWAS of self-reported sleep duration in 446,118 individuals from the UK Biobank, we identified 78 associated loci at *P*<5×10^-8^ (Dashti *et al.* BioRxiv 2018, https://doi.org/10.1101/274977). Sixty-seven (85.9%) of these SNPs were directionally consistent between the self-report and activity monitor derived sleep duration GWAS (*P*_*binomial*_ = 6×10^-11^; **Figure 3** and Dashti *et al.* BioRxiv 2018, https://doi.org/10.1101/274977). Furthermore, in a parallel report (Jones *et al.* BioRxiv 2018, https://doi.org/10.1101/303925) we have shown that of the 341 lead variants at self-reported chronotype loci, 310 (90.9%) had a consistent direction of effect for accelerometer-derived midpoint-sleep (*P*_*binomial*_ = 5×10^-59^), 316 (92.7%) with L5 timing (*P*_*binomial*_ = 3×10^-65^) and 310 (90.9%) with M10 timing (*P*_*binomial*_ = 5×10^-59^; Jones *et al.* BioRxiv 2018, https://doi.org/10.1101/303925). **Figure 4** shows a scatter plot of self-reported associated chronotype effects against L5 timing effects.

## DISCUSSION

Our analysis presents the first large-scale GWAS of multiple sleep traits estimated from accelerometer data using our validated activity-monitor sleep algorithm^28^,^29^. We have identified 47 genetic associations at *P*<5×10^-8^ across 7 traits representing sleep duration, quality and timing. These loci included 10 novel variants for sleep duration and 26 for sleep quality not detected in larger studies of self-reported sleep traits.

Of the novel associated loci, a low frequency (MAF=4.2%) missense variant (p.Tyr727Cys) at the *PDE11A* locus (rs17400325) was associated with sleep duration and sleep efficiency. The variant was associated with sleep duration (*P*=0.004) in the meta-analysis of the replication cohort. Fine-mapping provided a high probability (>90%) that this is the causal variant at the locus. This variant has previously been associated with migraine and near-sightedness in a scan of 42 traits from 23andMe^42^. In the UK Biobank the variant was not associated with migraine (*P*=0.44), consistent with the latest migraine meta-analysis where it was not amongst the associated loci^43^, but was associated with myopia (*P*=9×10^-10^). The allele which associates with reduced risk of myopia is associated with increased sleep efficiency and duration. Protein truncating variants in *PDE11A* have been suggested to cause adrenal hyperplasia^44^; however, one of these variants (R307X, rs76308115) is present at 0.5% frequency in the UK Biobank (with 11 rare allele homozygotes) and is not associated with sleep efficiency (P=0.99) or duration (*P*=0.54). This suggests that if Tyr727Cys *PDE11A* is the causal variant at this locus then it is an activating mutation. *PDE11A* is expressed in the hippocampus and it has been suggested as a potential biological target for interventions in neuropsychiatric disorders^45^.

Our analysis identified variants in loci that were enriched for genes involved in the serotonin pathway - the strongest pathway associated with sleep quality. Serotonergic transmission plays an important role in sleep cycles^46^,^47^. High levels of serotonin are associated with wakefulness and lower levels with sleep. Furthermore, serotonin is synthesized by the pineal gland as a processing step for melatonin production, a key hormone in circadian rhythm regulation and sleep timing. Melatonin is frequently taken as a dietary supplement in the United States with its use more than doubling between 2007 and 2012^48^, although clinical trial results for sleep and circadian rhythm disorders are mixed^49^. In addition, excess melatonin levels can also lead to disturbed sleeping and other health issues with the American Academy of Sleep Medicine recommending avoiding melatonin for chronic insomnia^50^.

A subset of variants previously associated with restless legs syndrome were associated with sleep duration, quality and timing measures. This observation is unlikely to be an artefact caused by limb movements during sleep because we found that the same variants are associated with self-report measures of sleep duration, chronotype and insomnia. Therefore, it seems likely that we are detecting how restless legs syndrome can influence sleep. In the UK Biobank, restless legs syndrome was only identified through the Hospital Episodes Statistics (HES) data using the ICD-10 code “G25.8” (“Other specified extrapyramidal and movement disorders”), the parent category of the more specific “G25.81” code (“Restless legs syndrome”). Under the assumption that all individuals reporting “G25.8” had restless legs, we observed 38 individuals within our accelerometer subset. Removing these individuals did not change our conclusions. Studies with more in-depth phenotyping of sleep disorders are needed to more fully evaluate the contribution of RLS to sleep traits.

Our Mendelian Randomization analysis also provides some evidence of a causative effect of higher waist-hip-ratio (adjusted for BMI) on lower sleep duration and lower sleep efficiency. This suggests that fat distribution plays a role in sleep, although there was also a nominal causative association with BMI which also suggests a general role of overall adiposity. We also observed evidence of a causative association between higher educational attainment and lower sleep duration. Both the adiposity and educational attainment MR results were robust to a range of MR sensitivity analyses (**Supplementary Table 12**). We did not observe evidence of a causal effect of accelerometer-derived sleep variables on genetically correlated traits. This may be due to the relatively limited power because of the relatively small number of genetic instruments available.

Our data provide strong evidence that some accelerometer-derived measures of sleep provide higher precision than self-report measures, whilst for others there is little gain through objective measurement with questionnaire data being just as effective. For example, of the 11 accelerometer-based sleep duration loci we identified, only one (the *PAX8* variant) had been previously identified in self-reported sleep duration GWAS despite these studies having much larger sample sizes. Variants with nominal evidence of association with self-reported sleep duration had weaker effects. This difference may be due to reporting biases related to the UK Biobank questionnaire (e.g. response was in hourly increments) and due to asking participants to include nap-time in their sleep duration. In contrast the accelerometer derived estimates of L5 timing, the least active 5 hours of the day, correlated well with self-report estimates. These data suggest that the answer to the very simple question “are you a morning or evening person” provides similar power as wearing accelerometers for 7 days and nights. In a parallel GWAS analysis, the *PAX8* variant was also associated with self-report insomnia (Lane *et al.* BioRxiv 2018: https://doi.org/10.1101/257956). In addition, five of the loci were nominally associated (*P*<0.05) with either self-report sleep-duration or insomnia. At least two of the sleep duration signals have been previously associated with mental health disorders including schizophrenia and migraine^42^,^51^.

The Alzheimer’s disease risk allele at the *APOE* locus was seen to have apparently paradoxical associations with sleep related traits. Given the well-established association between the ε4 allele and greater risk of Alzheimer’s disease, we would not expect associations between this allele and higher sleep quality considering previously observed associations of sleeping patterns with cognitive decline and Alzheimer’s disease^4^. A similar paradoxical association was also reported recently in a study of over 2,300 men aged over 65 with overnight PSG data that showed the total time in stage N3 sleep was higher for individuals carrying two copies of ε*4* compared with those carrying one or zero copies^52^. Furthermore, a recent genetic study of physical activity also identified a paradoxical association between the ε4 allele and increased levels of physical activity (Klimentidis *et al,* BioRxiv 2017, https://doi.org/10.1101/179317). The more likely explanations for these associations we suggest are ascertainment and survival bias. The UK Biobank participants ranged from 44 to 79 years of age when wearing the accelerometer devices. Older UK Biobank participants, with the highest risk of cognitive decline with an ε*4/*ε*4* haplotype and agreeing to an accelerometer-based experiment could be protected from cognitive decline because of selection bias due to other factors^53^. Consistent with this potential bias, the ε*4* allele association with reduced numbers of nocturnal sleep episodes is stronger in older age. For example, when splitting individuals by median age, the per allele effect on number of sleep episodes was twice that of the older versus younger group.

There are some limitations to this study. First, a sleep diary was not collected by the UK Biobank participants, a traditional tool to guide the start and end timing of nocturnal sleep episodes, commonly used in actigraphy studies. We have developed and used an open source method to overcome the lack of a sleep diary that has been validated against polysomnography^28^,^29^ to estimate sleep onset and waking up time. However, as no sleep diary data exists it is hard to define bedtime prior to sleep, resulting in the inability to characterise phenotypes such as sleep onset latency (the time between going to bed and falling asleep). Second, the activity monitors were worn up to 10 years from when baseline data was collected. Despite this, the correlation between self-report and activity measures of sleep duration was consistent with previous studies, and the correlation did not differ based on time between baseline (self-report time) and accelerometer wear when splitting by time-difference deciles (*r* = −0.03, *P* = 0.94). Third, due to relatively small sample sizes of replication studies, we had limited power to replicate associations identified in the UK Biobank. The variance explained by individual variants in the UK Biobank ranged from 0.03% to 0.19%, for which we had <63% power to detect at a statistical threshold of *P*=0.001 (accounting for 47 tests) in the meta-analysis of 4,401 individuals. However, we observed an enrichment for directional consistency in effect estimates in the replication meta-analysis and in combined-effects analyses identified associations for sleep duration, sleep efficiency, number of nocturnal sleep episodes and sleep timing. Finally, the UK Biobank participants are not representative of the UK population, as participants had a higher socio-economic status overall and were healthier, on average, given the prevalence of diseases amongst the participants^53^,^54^. This was particularly true of the participants who took part in the activity monitor study.

In conclusion, we have performed the first large-scale GWAS of objective sleep measures. We demonstrate that self-report measures are good proxies for objective sleep measures, but use of objectively measures of sleep quality allowed us to identify additional loci not identified by previous self-report GWAS studies including potential new therapeutic targets for poor sleep.

## METHODS

### Data availability

The full set of GWAS summary statistics for all eight accelerometer-based measures are available at http://www.t2diabetesgenes.org/data/.

### UK Biobank participants

The study population was drawn from the UK Biobank study – a longitudinal population-based study of individuals living in the UK^54^. Analyses were based on individuals estimated to be of European ancestry. European ancestry was defined through the projection of UK Biobank individuals into the principal component space of the 1000 Genomes Project samples^55^ and subsequent clustering based on a *K*- means approach, centering on the means of the first 4 principal components.

### Genetic Data

Imputed genetic data was downloaded from the UK Biobank (Bycroft, *et al.* BioRxiv 2017, https://doi.org/10.1101/166298). We limited our analysis to 11,977,111 genetic variants imputed using the Haplotype Reference Consortium imputation reference panel with a minimum minor allele frequency (MAF) > 0.1% and imputation quality score (INFO) > 0.3.

### Activity-monitor Devices

A triaxial accelerometer device (Axivity AX3) was worn between 2.8 and 9.7 years after study baseline by 103,711 individuals from the UK Biobank for a continuous period of up to 7 days. Details of data collection and processing have been previously described^56^. Of these 103,711 individuals, we excluded 11,067 individuals based on activity-monitor data quality. This included individuals flagged by UK Biobank as having data problems (field 90002), poor wear time (field 90015), poor calibration (field 90016), or unable to calibrate activity data on the device worn itself requiring the use of other data (field 90017). Individuals were also excluded if number of data recording errors (field 90182), interrupted recording periods (field 90180), or duration of interrupted recoding periods (field 90181) was greater than the respective variable’s 3^rd^ quartile + 1.5×IQR. Phenotypes determined using the SPT-window (all phenotypes except L5 and M10 timing) had additional exclusions based on short (<3 hours) and long (>12 hours) mean sleep duration and too low (<5) or too high (>30) mean number of sleep episodes per night (see below). These additional exclusions were to ensure that individuals with extreme (outlying), and likely incorrect, sleep characteristics were not included in any subsequent analyses. A maximum of 85,670 individuals remained for our analyses.

### Accelerometer data processing and sleep measure derivations

We derived 8 measures of sleep quality, quantity and timing. All measures were derived by processing raw accelerometer data (.cwa). We first converted the .cwa files available from the UK Biobank to .wav files using “omconvert” for signal calibration to gravitational acceleration^56^,^57^ and interpolation^56^. The .wav files were processed with the open source R package GGIR^29^ (http://doi.org/10.5281/zenodo.1175883 (Version v1.5-17)) to infer accelerometer non-wear time^58^, and extract the z-angle across 5-second epochs from the time-series data for subsequent use in estimating the sleep period time window^29^ and sleep episodes within it^28^.

*Sleep period time window (SPT-window).* The SPT-window was estimated using a validated algorithm previously described^29^ and implemented in the GGIR R package. Briefly, for each individual, median values of the absolute change in estimated z-angle (representing the dorsal-ventral direction when the wrist is in the anatomical position) across 5-minute rolling windows were calculated across a 24-hour period, chosen to make the algorithm insensitive to accelerometer orientation. The 10^th^ percentile was incorporated into the threshold distinguishing movement from non-movement. Bouts of inactivity lasting ≥30 minutes are recorded as inactivity bouts. Inactivity bouts that are <60 minutes apart are combined to form inactivity blocks. The start and end of the longest block defined the start and end of the SPT-window.

*Sleep duration and variability*. Sleep episodes within the SPT-window were defined as periods of at least 5 minutes with no change larger than 5° associated with the z-axis of the activity-monitor, as motivated and described in van Hees *et al.* (2015). The summed duration of all sleep episodes was used as indicator of sleep duration within the SPT-window. The total duration over the activity-monitor wear-time was averaged. Individuals with an average sleep duration <3 hours or >12 hours were excluded from all analyses. In addition, the standard deviation of sleep duration was also calculated and put forward for statistical analysis for individuals with 7 days of accelerometer wear.

*Sleep efficiency.* This was calculated as sleep duration (defined above) divided by the time elapsed between the start of the first inactivity bout and the end of the last inactivity bout (which equals the SPT-window duration).

#### Number of nocturnal sleep episodes within the SPT-window

This was defined as the number of sleep episodes within the SPT-window. Individuals with an average number of nocturnal sleep episodes ≤5 or ≥30 were excluded from all analyses.

*Least active 5 hours (L5) timing.* The mid-point of the least-active 5 hours (L5) of each day were defined as the 5-hour period with the minimum average acceleration. These periods were estimated using a rolling 5-hour time window. The midpoint was defined as the number of hours elapsed since the previous midnight (for example, 7pm = 19 and 2am = 26). Days with <16 hours of valid-wear time (as estimated by GGIR) were excluded from L5 estimates.

*Most-active 10 hours (M10) timing.* The mid-point of the most-active 10 hours (M10) of each day were defined as the 10-hour period with the maximum average acceleration. These periods were estimated using a rolling 10-hour time window. The midpoint was defined as the number of hours elapsed since the previous midnight. Days with <16 hours of valid-wear time (as estimated by GGIR) were excluded from M10 estimates.

*Sleep-midpoint timing.* Sleep midpoint was calculated for each sleep period as the midpoint between the start of the first detected sleep episode and the end of the last sleep episode used to define the overall SPT-window (above). This variable is represented as the number of hours from the previous midnight.

*Diurnal inactivity.* Diurnal inactivity was estimated by the total daily duration of estimated bouts of inactivity that fell outside of the SPT-window. This measure captures very inactive states such as napping and wakeful rest but not inactivity such as sitting and reading or watching television, which are associated with a low but detectable level of movement.

### Comparison against self-reported sleep measures

We performed analyses of self-reported measures of sleep. Self-reported measures analysed included a) the number of hours spent sleeping over a 24-hour period (including naps); b) insomnia; c) chronotype – where “definitely a ‘morning’ person”, “more a ‘morning’ than ‘evening’ person”, “more an ‘evening’ than a ‘morning’ person”, “definitely an ‘evening’ person” and “do not know”, were coded as 2, 1, −1, −2 and 0 respectively, in our continuous variable.

### Statistical Analysis

*Genome-wide association analyses*. We performed all association tests in the UK Biobank using BOLT-LMM v2.3^59^, which applies a linear mixed model (LMM) to adjust for the effects of population structure and individual relatedness, and enables the inclusion of all related individuals in our white European subset, boosting our power to detect associations. This meant a sample size of up to 85,670 individuals, as opposed to a maximal set of 72,696 unrelated individuals. Prior to association testing, phenotypes were first adjusted for age at accelerometry, sex, study centre, and season when activity monitor worn (categorical). All phenotypes except sleep duration variation were also adjusted for the number of measurements used to calculate each participant’s measure (number of L5/M10 measures for L5/M10 timing, number of days for diurnal inactivity and number of nights for all other phenotypes). At runtime, association tests included genotyping array (categorical; UKBileve array, UKB Axiom array interim release and UKB Axiom array full release) as a covariate.

*SNP-based heritability analysis.* We estimated the pseudo-heritability of the eight derived accelerometer traits using BOLT-REML (version 2.3.1)^59^. We used 524,307 high-quality genotyped single nucleotide polymorphisms (SNPs) (bi-allelic; MAF ≥1%; HWE *P*>1×10^-6^; non-missing in all genotype batches, total missingness <1.5% and not in a region of long-range LD^60^) to build the relatedness model and thus to estimate heritability. For LD structure information, we used the default 1000 Genomes ‘LD-Score’ table provided with the BOLT-REML software.

*Gene-set, tissue expression enrichment, and overlap with GWAS-catalog analyses.* Gene-set analyses and tissue expression analyses were performed using MAGMA^40^ as implemented in the online Functional Mapping and Annotation of Genome-Wide Association Studies (FUMA) tool^61^. Analysis of differentially expressed genes was based on data from GTEx v6 RNA-seq data^62^. Enrichment analyses of the overlap with associations previously reported through GWAS was also implemented through FUMA. Enrichment P-values for the proportion of overlapping genes present was based on the NIH GWAS catalog^63^.

#### Fine-mapping association signals

Fine-mapping analyses were performed using FINEMAP v1.2^37^ using the software’s shotgun stochastic search function and by setting the maximum number of causal SNPs at each locus to 20. At each locus, we included only those with *P*<0.01 and within 500Kb either side of the index variant to limit the number of SNPs in the analysis. We constructed the LD matrix by calculating the Pearson correlation coefficient for all SNP-SNP pairs using SNP dosages derived from the unrelated European subset of the full UK Biobank imputed genotype probabilities (N=379,769). We considered a SNP to be causal if it’s log_10_ Bayes factor was greater than 2, as recommended in the FINEMAP manual (http://www.christianbenner.com/index_v1.2.html).

#### Alamut annotation and eQTL mapping

We performed variant annotation of our fine-mapped loci using Alamut Batch v1.8 (Interactive Biosoftware, Rouen, France) using all default options and genome assembly GRCh37. For each annotated variant, we retained only the canonical (longest) transcript and reported the variant location, coding effect and the predicted local splice site effect. To investigate whether the fine-mapped SNPs were eQTLs, we searched for our SNPs in the single-tissue cis-eQTL dataset (v7), available at the GTEx portal (https://www.gtexportal.org/home/datasets) for significant SNP-gene eQTL associations. We reported a SNP as an eQTL for a gene if the SNP-gene association was significant for at least one tissue.

*Replication of findings.* Associations reaching *P*<5×10^-8^ were followed up in the CoLaus, Whitehall and Rotterdam studies. The GENEActiv accelerometer was used by the CoLaus and Whitehall studies and worn on the wrist by the participants. In the CoLaus study, 2,967 individuals wore the accelerometer for up to 14 days. Of these, 10 were excluded because of insufficient data, 234 excluded as non-European, and a further 148 were excluded due to an average sleep duration of less than 3 hours or more than 12 hours. A total of 2,575 individuals remained for analysis of which 2,257 had genetic data. In the Whitehall study, 2,144 were available for analysis, with the GENEActiv accelerometer worn for up to 7 days having performed the same exclusions. The Rotterdam Study used the Actiwatch AW4 accelerometer device (Cambridge Technology Ltd.). Genetic association analysis was based on imputed data (where available) and performed using standard multiple linear regression. Overall summary statistics were obtained through inverse-variance based meta-analysis implemented in METAL^64^. Combined variant effects on respective traits were subsequently calculated using the ‘metan’ function in STATA using the betas and standard errors obtained through the primary meta-analysis of the three replication studies.

*Sensitivity Analysis*. To assess whether stratification was responsible for any of the individual variant associations in a subset of the cohort, we performed multiple sensitivity analyses in unrelated European subsets of the UK Biobank using STATA. The sensitivity analyses carried out were: 1) males only, 2) females only 3) individuals younger than the median age (at start of the activity monitor wear period), 4) individuals older than the median age, 5) adjustment for body mass index (BMI) (UK Biobank data field 21001), 6) adjusting for BMI and lifestyle factors and 7) excluding individuals working shifts, taking medication for sleep or psychiatric disorders, self-reporting a mental health or sleep disorder, or diagnosed with depression, schizophrenia, bipolar disorder, anxiety disorders or mood disorder in the HES data (see **Supplementary Methods**). The sensitivity analyses were performed by regressing the phenotype against the variant dosage, adjusting for the same covariates as described for the BOLT-LMM GWAS and additionally adjusting for the first 5 principal components to account for population structure. All exclusions and adjustments were made using baseline records (taken at the assessment centre).

*Mendelian Randomisation (MR)*. We performed two-sample MR, using the inverse variance weighted approach^65^ as our main analysis method, and MR-Egger^65^, weighted median estimation^66^ and penalised weighted median estimation^66^ as sensitivity analyses in the event of unidentified pleiotropy of our genetic instruments. MR results may be biased by horizontal pleiotropy, i.e. where the genetic variants that are robustly related to the exposure of interest independently influence the outcome, through association with another risk factor for the outcome. IVW assumes that there is either no horizontal pleiotropy (under a fixed effect model) or, if implemented under a random effects model after detecting heterogeneity amongst the causal estimates, that (i) the strength of association of the genetic instruments with the risk factor is not correlated with the magnitude of the pleiotropic effects, and (ii) the pleiotropic effects have an average value of zero. MR-Egger provides unbiased causal estimates if just the first condition above holds, by estimating and adjusting for non-zero mean pleiotropy. The weighted median approach is valid if less than 50% of the weight in the analysis stems from variants that are pleiotropic (i.e. no single SNP that contributes 50% of the weight or a number of SNPs that together contribute 50% should be invalid because of horizontal pleiotropy). Given these different assumptions, if all methods are broadly consistent, our causal inference is strengthened.

In an effort to reduce the number of genetic instruments violating the above assumptions, we used a newly-described method (Bowden *et al.* BioRxiv 2017, http://dx.doi.org/10.1101/159442) to quantify, using a new iterative weighting method, each instrument’s contribution to heterogeneity of the causal IVW estimate. High heterogeneity in Cochran’s Q statistic, which should follow a 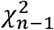 distribution for n instruments, indicates that either invalid (horizontally-pleiotropic) instruments have been included or that MR modelling assumptions have been violated. We therefore excluded variants with an extreme Cochran’s Q greater than the Bonferroni corrected threshold (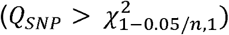) prior to performing MR analysis.

## ACKNOWLEDGMENTS

This research has been conducted using the UK Biobank Resource (application 9072). S.E.J. is funded by the Medical Research Council (grant: MR/M005070/1). S. Sabia is supported by EC Horizon2020 (LIFEPATH 633666). J.T. is funded by a Diabetes Research and Wellness Foundation Fellowship. M.A.T., M.N.W. and A.M. are supported by the Wellcome Trust Institutional Strategic Support Award (WT097835MF). A.R.W., T.M.F and H.Y. are supported by the European Research Council grants: SZ-245 50371-GLUCOSEGENES-FP7-IDEAS-ERC and 323195. R.N.B. and R.M.F. are funded by the Wellcome Trust and Royal Society, grant 104150/Z/14/Z. The generation and management of GWAS genotype data for the Rotterdam Study (RS I, RS II, RS III) was executed by the Human Genotyping Facility of the Genetic Laboratory of the Department of Internal Medicine, Erasmus MC, Rotterdam, The Netherlands. The GWAS datasets are supported by the Netherlands Organisation of Scientific Research NWO Investments (nr. 175.010.2005.011, 911-03-012), the Genetic Laboratory of the Department of Internal Medicine, Erasmus MC, the Research Institute for Diseases in the Elderly (014-93-015; RIDE2), the Netherlands Genomics Initiative (NGI)/Netherlands Organisation for Scientific Research (NWO) Netherlands Consortium for Healthy Aging (NCHA), project nr. 050-060-810. We thank Pascal Arp, Mila Jhamai, Marijn Verkerk, Lizbeth Herrera and Marjolein Peters, MSc, and Carolina Medina-Gomez, MSc, for their help in creating the GWAS database, and Karol Estrada, PhD, Yurii Aulchenko, PhD, and Carolina Medina-Gomez, MSc, for the creation and analysis of imputed data. The Rotterdam Study is funded by Erasmus Medical Center and Erasmus University, Rotterdam, Netherlands Organization for the Health Research and Development (ZonMw), the Research Institute for Diseases in the Elderly (RIDE), the Ministry of Education, Culture and Science, the Ministry for Health, Welfare and Sports, the European Commission (DG XII), and the Municipality of Rotterdam. The authors are grateful to the study participants, the staff from the Rotterdam Study and the participating general practitioners and pharmacists. The Rotterdam Study has been approved by the Medical Ethics Committee of the Erasmus MC (registration number MEC 02.1015) and by the Dutch Ministry of Health, Welfare and Sport (Population Screening Act WBO, license number 1071272-159521-PG). The Rotterdam Study has been entered into the Netherlands National Trial Register (NTR; www.trialregister.nl) and into the WHO International Clinical Trials Registry Platform (ICTRP; www.who.int/ictrp/network/primary/en/) under shared catalogue number NTR6831. The CoLaus study was and is supported by research grants from GlaxoSmithKline, the Faculty of Biology and Medicine of Lausanne, and the Swiss National Science Foundation (grants 33CSCO-122661, 33CS30-139468 and 33CS30-148401). The institutional Ethics Committee of the University of Lausanne, which afterwards became the Ethics Commission of Canton Vaud (www.cer-vd.ch) approved the baseline CoLaus study (reference 16/03, decisions of 13th January and 10th February 2003). The approval was renewed for the first (reference 33/09, decision of 23rd February 2009), the second (reference 26/14, decision of 11th March 2014) and the third (reference PB_2018-00040, decision of 20th March 2018) follow-ups. The Whitehall II study is supported by grants from the US National Institutes on Aging (R01AG013196; R01AG034454), the UK Medical Research Council (MRC K013351; MR/R024227/1), and British Heart Foundation (RG/13/2/30098). The University College London Hospital Committee on the Ethics of Human Research approved the study (reference number 85/0938). All participants provided written informed consent to participate in the study and to have their information obtained from treating physicians.

## AUTHOR CONTRIBUTIONS

The study was designed by T.M.F., P.G., V.T.v.H., S.E.J., D.R.M., M.N.W. and A.R.W. Participation in acquisition, analysis and/or interpretation of data was performed by N.A., R.N.B., H.S.D., J.E., T.M.F., R.M.F., J.W.H., V.T.v.H., Y.J., S.E.J., D.K., M.Z., Z.K., A.I.L., D.R.M., A.M., K.S.R., S.Sabia, R.S., S.Sharp, A.v.d.S., H.T., M.A.T., J.T., P.M.V., M.N.W., A.R.W. and H.Y. Main writing group comprised of H.S.D. T.M.F., P.G., V.T.v.H, S.E.J., D.K., J.M.L., A.I.L. D.R.M., S.Sabia, H.T., M.K.T., P.M.V., M.N.W. and A.R.W. All authors reviewed this manuscript. A.R.W. is the guarantor of this work and, as such, had full access to all the data in the study and takes responsibility for the integrity of the data and the accuracy of the data analysis.

## Supplementary Tables

**Supplementary Table 1.** Descriptive statistics of sleep and activity measures derived from accelerometer data. Units for L5 timing, M10 timing, sleep duration, sleep duration variation (SD), sleep midpoint and diurnal inactivity are in hours. Sleep efficiency is a ratio and number of sleep episodes is a count.

**Supplementary Table 2:** Spearman’s rank correlation statistics for activity monitor derived sleep traits and self-report hours slept and self-report chronotype (coded for increased morningness).

**Supplementary Table 3.** Association statistics for the 47 signals discovered in UKB Biobank in the Whitehall and CoLaus and Rotterdam I, II, and III replication cohorts. Meta-analysis of the results across the studies are also provided. Grey cells indicate data unavailable from particular study.

**Supplementary Table 4.** Analysis of the combined genetic effects from Supplementary Table 3 within the Whitehall, CoLaus and Rotterdam replication studies.

**Supplementary Table 5.** Averages of the mean number of nocturnal sleep episodes detected within individuals in the UK Biobank split by APOE Alzheimer’s disease risk haplotypes and lower/upper age groups.

**Supplementary Table 6.** Results from sensitivity analyses performed for the 47 signals reaching *P*<5×10^-8^ in the UK Biobank. All analyses were performed in a set of unrelated European where individuals from related pairs were removed at random. Association tests were carried out for all phenotypes on both the raw scale and inverse-normalised scale. Sensitivity analysis included: 1) in males only, 2) in females only, 3) in those lower than median age at actigraphy (63.7 years), 4) in those greater than or equal to the median age, 5) in all European unrelated but adjusting for BMI in addition to standard adjustments, 6) in all European unrelated but also adjusting for BMI and lifestyle factors, and 7) excluding those reporting shift work, having self-report or hospital-recorded mental health or sleep disorders, and those taking anxiolytic, antipsychotic, antidepressant or sleep medication. Lifestyle adjustments for analysis (6) and exclusions for analysis (7) are described in greater detail in the Supplementary Methods.

**Supplementary Table 7.** Association result cross-tabulation against other traits for the 47 SNPs representing genetic associations reaching *P*<5×10^-8^ in UK Biobank. Cross-tabulation also includes results based on the latest self-report chronotype meta-analyses (Jones *et al.*, BioRxiv 2018, https://doi.org/10.1101/303925), self-report Insomnia GWAS (Lane et al., BioRxiv 2018, https://doi.org/10.1101/257956) and sleep duration GWAS in UK Biobank are also provided.

**Supplementary Table 8.** Fine-mapped loci with at least one “plausible” variant (log10 Bayes’ Factor > 2) with variant annotations from GTEx and Alamut.

**Supplementary Table 9.** MAGMA Gene-Set Analysis for SNPs associated with disturbed sleep based on number of nocturnal sleep episodes reaching Bonferroni significance.

**Supplementary Table 10.** Results from Mendelian Randomization (MR) analyses of Restless Legs Syndrome exposure against multiple outcomes using 4 methods: 1) using Inverse-variance (IV) weighted MR, 2) Egger MR, 3) Weighted Median (WM) MR. 4) Penalised-weighted mean (PWM).

**Supplementary Table 11.** Genetic correlation results for the 8 accelerometer-derived sleep traits against 234 LD Hub phenotypes, ordered by *P*-value. *P*-values reaching Bonferroni significance (*P*<0.05/(8*234)) in bold.

**Supplementary Table 12.** Mendelian Randomization analyses testing causality of seven genetically correlated traits on accelerometer-based sleep outcomes.

**Supplementary Table 13.** Mendelian Randomization analyses testing causality of four accelerometer-based sleep exposures on genetically correlated traits. Sleep exposures with <3 genome-wide associations at *P*<5×10^-8^ or with <3 genetic instruments available in published datasets (highlighted grey) were excluded from this analysis.

